# Secondary lymphoid organ stroma activate and elaborate regulatory T cells to suppress autoantibody production in a novel model of systemic autoimmunity

**DOI:** 10.1101/2025.04.21.649586

**Authors:** Tia Y. Brodeur, Ann Marshak-Rothstein

## Abstract

Preclinical models of lupus indicate that T cell-B cell collaboration drives pathogenic antinuclear antibody (ANA) production and sustains T cell activation. Mechanisms that normally limit T cell activation of autoreactive B cells remain incompletely resolved but potentially include the absence of autoreactive effector T cell subsets and/or the presence of autoantigen-specific regulatory T cells (Tregs). Several studies have addressed this issue by using experimental systems dependent on transgenic autoreactive B cells, but much less is known about the activation of autoreactive B cells present in a polyclonal repertoire. We have now explored the role of effector T cells and Tregs using mice that express an inducible pseudo-autoantigen on antigen presenting cells (APCs) including a normal B cell repertoire. Bone marrow chimera experiments demonstrated that radioresistant non-hematopoietic stromal APCs (rAPC) present in the thymus and in peripheral lymphoid tissue induced the differentiation and expansion of a subset of CD62L^+^CD69^+^ Tregs associated with decreased autoantibody production and MHC-II. In this study, we show that secondary lymph node stromal cells may be crucial for suppression of endogenous autoreactive B cells in the setting of robust T cell help.

## Introduction

Systemic lupus erythematosus (SLE) is a progressive, multisystem autoimmune disease characterized by anti-nuclear autoantibody (ANA) production by aberrantly activated autoreactive B cells and immune complex deposition in the kidneys, joints, vasculature, and skin. SLE is also characterized by elevated inflammatory cytokine levels in the serum and persistent autoreactive T cell activity. T and B cells are thought to have a cooperative relationship in SLE, as B cell deficiency significantly reduces the number of T cells exhibiting an activated/memory phenotype and subsequent T cell infiltration of the kidneys in autoimmune prone mice (1-3). Moreover, B cell depletion therapy, including use of the anti-CD20 antibody rituximab, can lead to the loss of activated T cells in lupus patients and decreased ANA titers (4, 5). Consequently, the biology of autoreactive B cells is an area of active investigation.

Autoreactive B cells exist in healthy humans and mice and can persist as fully reactive “ignorant” cells or in variable states of anergy. Although anergic autoreactive B cells often respond poorly to B cell receptor (BCR) engagement alone, they can be rescued when provided with T cell help. For example, anti-DNA B cells (3H9) are anergic, but can be re-activated by provision of cognate interactions with T cells (6). Prior studies also point to a role for autoreactive T cells in the activation of ignorant B cells (7). The contribution of effector T cells to autoantibody production has also been studied *in vivo*. Strategies include the depletion of CD4^+^ T cells from autoimmune prone mice (8-11) and the use of mice genetically deficient for key T cell cytokines such as IL-4 and IFN-γ (12, 13). In most murine models, Th1 cells, which produce IFN-γ and IL-2, have been identified as pathogenic (14) and a Th1 signature has been identified in SLE patients (15). Th2 cells are effective B-helper cells at least in part through production of IL-4, and have been largely understudied in the context of lupus despite the fact that IgE autoantibodies— a Th2-dependent isotype—are increased in patients with active SLE (16). Thus, the contribution of Th2 cells to SLE warrants further investigation.

Treg development in the thymus depends in part on non-hematopoietically derived radioresistant cells in the thymus, namely medullary and cortical thymic epithelial cells (mTECs and cTECs). mTECs present self-peptide:MHC-II complexes directly and mTEC-derived self-antigen (Ag) is presented by neighboring dendritic cells (DCs) (17, 18). As in the thymus, non-hematopoietic APCs in secondary lymphoid organs (SLO) can express transcription factors such as AIRE and Deaf1, which drive ectopic expression of tissue-specific Ags. These lymph node stromal cells (LNSC) have an important role in preventing autoimmunity by both constitutively expressing and presenting tissue-specific Ags and decreasing T cell proliferation. Specifically, LNSC mediate CD8^+^ T cell tolerance through the deletion of autoreactive CD8^+^ T cells and iNOS-dependent suppression of proliferation (19-23). The role of LNSC in regulating CD4+ T cell tolerance—especially in relation to Tregs—is less clear, however; LNSC constitutively express low levels of MHC-II and upregulate MHC-II in response to inflammatory stimuli(24). Additionally, the interaction between LNSC and Treg regulates tissue-specific immunity in a transplant model (25).

We have developed a model to study cognate interactions between autoreactive T cells and self-Ag reactive B cells in the presence or absence of LNSC presentation of self-Ag. In this manuscript, we show that T cell activation is an important checkpoint in the prevention of autoantibody production, that autoreactive T cell activation is sufficient for autoantibody production by autoreactive B cells, and that stromal self-Ag presentation is required to most effectively control autoantibody responses.

## Results

### Development of a transgenic mouse expressing an inducible autoantigen

To study autoreactive B cell responses in the presence or absence of cognate help from activated autoreactive T cells, we generated double transgenic mice that could be induced to express a tissue-restricted pseudo-autoantigen. One transgene incorporated a fusion protein encoding the **t**ransmembrane domain of **t**ransferrin (to target protein expression to the cell surface) and chicken **<u>o</u>**valbumin (the selected “pseudo-autoantigen” that elicits T cell activation) (26). This TGO construct was cloned into a tetracycline-responsive element (TRE)-containing plasmid and used to generate the TRE-TGO mouse. These mice can be crossed to strains that express a transgene encoding the reverse tetracycline transactivator (rtTA) under control of a tissue-specific promoter to allow TGO expression in response to tetracycline or the synthetic derivative doxycycline (Dox) (27). TRE-TGO mice were crossed to an Ii-rtTA transgenic line (28) to generate mice that could be induced to express OVA in all MHC class II^+^ cells (Supplemental Figure 1). This cross, referred to as Ii-TGO, enables TGO to serve as a pseudo-autoantigen where OVA can be presented to OVA-specific CD4^+^ T cells for as long as the mice are maintained on Dox (29) We previously confirmed inducible expression of TGO and antigen-dependent proliferation of OVA-specific transgenic DO11.10 (DO11) CD4^+^ T cells and subsequent tissue-specific autoimmunity (29).

### Ii-TGO mice develop lupus-like autoimmune disease following adoptive transfer of activated DO11.10 T cells

Chronic amplification loops due to sustained T cell-B cell interaction are thought to promote systemic autoimmunity in B cell-driven diseases such as lupus. For autoreactive Tg B cells activated by autoantigen complexes that incorporate endogenous TLR ligands, unprimed T cells have clearly been shown to enhance the extent of autoantibody production and isotype switching (6, 7). To determine whether DO11 T cells could drive autoantibody production in Ii-TGO B mice, we adoptively transferred either naïve DO11 cells or *in vitro* activated Th2 DO11 T cells (Supplemental Figure 1B) into Ii-TGO mice fed normal or Dox chow. Notably, neither the naïve DO11 nor activated T cells induced ANA production over the following 12 weeks, suggesting that infusion of naïve or activated autoreactive T cells alone was not sufficient to break tolerance. However, when Dox-fed recipient mice were irradiated (400R) and then injected with naïve or activated T cells, as early as 6 weeks after Th2 transfer, Ii-TGO mice produced autoantibodies with a predominantly antinuclear staining pattern (Supplemental Figure 1B). Importantly, when comparing total Ig isotype titers of the ANA+ recipients, IgG2a and IgG2b only increased two-fold (Supplemental Figure 1C), suggesting that autoreactive B cells were preferentially activated by DO11 Th2 cells over other (non-autoreactive) B cells in the peripheral repertoire.

### Ectopic expression of self-Ag on radioresistant APCs drives extensive Treg development independent of endogenous α-chain expression

At least two studies have shown that autoantibody production is dampened in mice immunized with self-Ag and adjuvant for a tissue-specific transgene-encode Ag if antigen-specific Tregs are present (30, 31). Similarly, Tregs can decrease activation of anti-dsDNA B cells in response to T cell help (6). Moreover, in previous studies, an unexpectedly high frequency of Treg cells were found in the thymus and skin draining lymph nodes of non-induced TGO mice crossed to DO11 mice, consistent with low level thymic TGO expression even in the absence of Dox (27) (Figure 1A).

**Figure 1.**
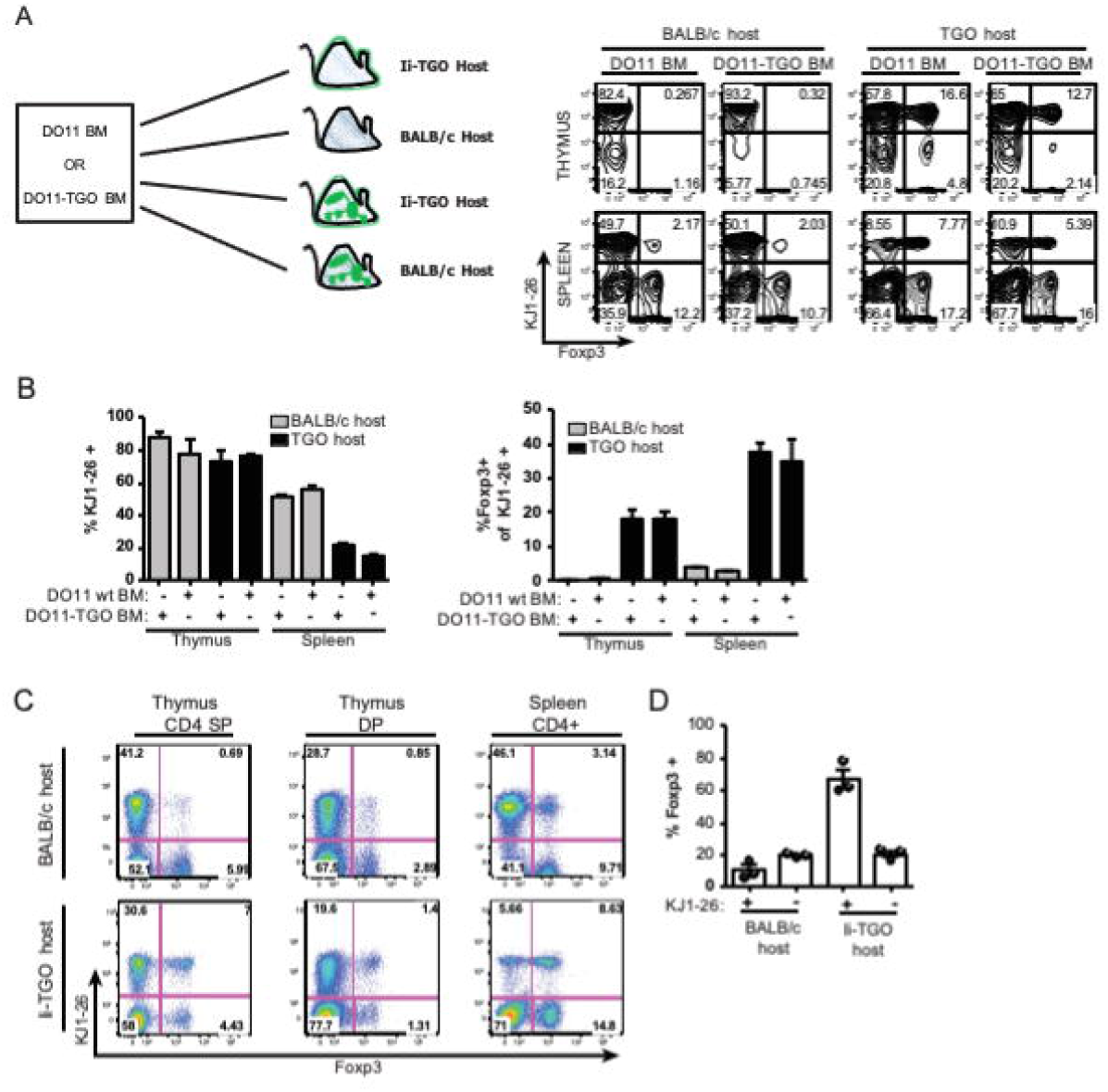
Ectopic expression of self-Ag drives extensive Treg development. (A) Bone marrow chimeras were generated using lethally irradiated BALB/c or TGO hosts reconstituted with DO11 or DO11 × TRE-TGO bone marrow. At 12 weeks, thymi and spleens were stained for CD4, CD8, DO11 TCR (KJ1-26 antibody), CD25, and Foxp3; cells were first gated on CD4 SP events. Data are representative of two experiments with 3–4 mice per group, and summarized in (B). The graph was generated from one experiment with three mice per group, and is representative of two experiments. Grey bars represent BALB/c hosts and black bars represent TGO hosts. (C) Radiation BM chimeras using 50% DO11 *Rag2-/-* BM and 50% BALB/c BM to reconstitute lethally irradiated BALB/c or Ii-TGO recipients. Thymus and spleen were analyzed by flow cytometry. These data are summarized in (D) Error bar shows SEM.

After confirming that basal TGO expression in DO11 × Ii-TGO mice resulted in a high frequency of DO11 Tregs and that induced TGO expression could drive DO11 Treg expansion, we considered the possibility that Tregs could limit autoreactive B cell activation in unirradiated Ii-TGO mice. To determine whether radioresistant stromal cells were responsible for ectopic expression and presentation of TGO, we generated bone marrow (BM) chimeras by using T-depleted bone marrow cells obtained from either DO11 mice or [DO11 × TRE-TGO] F1 mice to reconstitute lethally irradiated BALB/c or TRE-TGO recipients. In the BALB/c hosts, most (>85%) thymic CD4 single positive (SP) T cells expressed the DO11 TCR, regardless of whether the donor BM was DO11 or DO11 × TRE-TGO, and only a small frequency (<0.5%) of these cells expressed Foxp3. In contrast, in the TGO hosts, again most of the CD4 SP thymocytes were KJ126+, but as a result of host TGO expression, 15-22% became Foxp3+. In the spleen, approximately 50% of the donor derived T cells in both the DO11à BALB/c and DO11xTGO à BALB/c chimeras were KJ1-26^+^. However, very few of these KJ126^+^ cells were FoxP3+ Tregs, even though a relatively normal proportion of the clonotype negative T cells were Tregs. In contrast, in both the DO11à Tre-TGO and DO11xTGO à Tre-TGO chimeras, a significant proportion of the DO11 T cells were deleted and only ∼20% of the CD4 splenic T cells were KJ1-26+. Moreover, of the remaining KJ126+ cells, approximately 30-40% expressed Foxp3 (Figure 1A-B).

Although Treg enrichment in TCR Tg in cognate Ag-expressing hosts has been shown (27), it has been presumed that Treg development required a slightly higher affinity receptor conferred by endogenous α-chain expression. To determine whether DO11 cells could populate the Treg compartment without the capacity to express an endogenous TCR-α chain, we generated radiation BM chimeras using 50% DO11 *Rag2-/-* BM and 50% BALB/c BM to reconstitute lethally irradiated BALB/c or Ii-TGO recipients. In these mice, we could compare the frequency of Tregs in the normal BALB/c polyclonal CD4^+^ T cell repertoire and highly restricted DO11 T cells. In BALB/c hosts, roughly half of the CD4+ T cell population expressed the DO11 TCR, and 6–12% of DO11 T cells co-expressed CD25 and Foxp3. Surprisingly, in TGO hosts, most *Rag2-/-* DO11 cells found in the spleen were Tregs (Figure 1C-D). Our data show that basal levels of TGO expression increase DO11 TCR/self-Ag interactions to promote autoreactive CD4^+^ cell deletion and also drive Treg development.

### Absence of self-Ag-bearing stromal cells is permissive to persistent autoreactive T cell activation and enhanced autoreactive B cell activity

After determining that in our system, radioresistant cell-derived Ag was solely responsible for Treg generation, we next wanted to determine whether the magnitude or specificity of B cell ANA production could be constrained by Ag-specific Tregs. To this end, we generated BM chimeras with negligible numbers of TGO-specific Tregs by reconstituting lethally irradiated non-Tg BALB/c mice with Ii-TGO BM stem cells [Ii-TGOàBALB/ chimeras]. We also reconstituted lethally irradiated Ii-TGO mice with Ii-TGO stem cells [Ii-TGOàIi-TGO chimeras]. Twelve weeks after reconstitution, chimeric mice were placed on Dox to induce TGO expression peripherally, but they were not sublethally irradiated as in the experiment described in Supplemental Figure 1B. The chimeric mice were then injected with either naïve DO11 cells, DO11 Th2 cells, or left uninjected. Six weeks later, serum samples collected from the mice were assayed for the presence of ANAs. ANAs only developed in mice adoptively transferred with DO11 Th2 cells, suggesting that self-reactive (DO11) naïve T cell activation was effectively controlled even in the absence of self-Ag specific Tregs. However, the Ii-TGOàBALB/c chimera hosts developed higher ANA titers than the Ii-TGOàIi-TGO chimera hosts (Figure 2A). Surprisingly, the frequency of activated CD44^+^ CD69^+^ CD4+ T cells was similar in both Th2-injected Ii-TGOàBALB/c hosts and Th2-injected Ii-TGOàIi-TGO hosts (Figure 2B), while B cell CD86 and MHC-II were markedly elevated only in the BALB/c hosts (Figure 2C).

**Figure 2.**
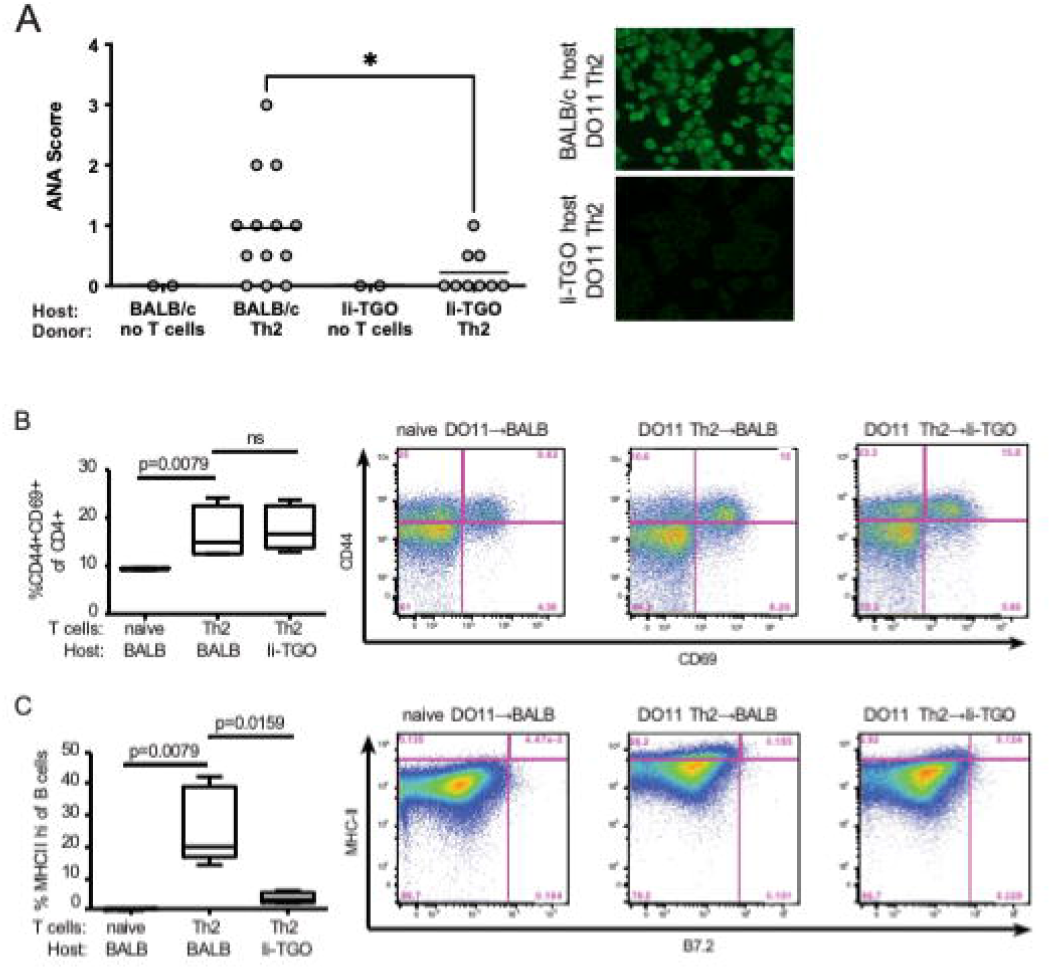
Absence of self-Ag bearing stromal cells is permissive to ANA production by autoreactive B cells without sublethal irradiation. Ii-TGOàBALB/c and Ii-TGOàIiTGO radiation chimeras were generated; 12-weeks post-irradiation, they were fed Dox chow and injected with PBS, naïve, or Th2-skewed DO11 T cells. (A) Six-weeks later, sera collected from these mice were analyzed for autoantibody production using a HEp2 immunofluorescent assay. Blinded **scoring was performed using a 0-4 scale relative to BALB/c-Fas**^**lpr/lpr**^ **serum. Combined data from two experiments are shown. A representative image to reflect the staining pattern is shown to the right of the graph. n for uninjected controls = 2, n for BALB/c host = 13, n for Ii-TGO host = 9. Unpaired *t*-test** was used for statistical analysis. (B, C) T cells were stained for markers of antigen experience/activation and analyzed by flow cytometry. B cells were stained for MHC-II. The gate for MHC-II^hi^ cells was set against a normal BALB/c control spleen. Box plots show five mice per group and are representative of two experiments. 2-way ANOVA was used for statistical analysis.

### Activation and elaboration of antigen-specific secondary lymphoid organ-resident Tregs are preferentially activated by radioresistant antigen-presenting cells

The abovementioned findings suggested a role for self-Ag presentation in the thymus in the control of aberrant ANA production by B cells. Because B cell activation and ANA production were reduced in the presence of stromal TGO, we sought to determine whether Treg activity was enhanced by peripheral expression of TGO to in some way facilitate Treg suppression of autoreactive B cells. To address this question, we generated BM chimeras by using DO11 x Ii-TGO F1 BM stem cells to reconstitute lethally irradiated BALB/c or Ii-TGO hosts. Twelve weeks after reconstitution, the mice were either kept on normal chow or placed on Dox chow for 7 days. Consistent with our previous experiments (Figure 2), in both the normal and Dox chow groups, the frequency and number of KJ126^+^ DO11 cells was decreased in Ii-TGO hosts compared to BALB/c hosts, and a comparably large fraction of DO11 cells co-expressed the Treg markers CD25 and Foxp3. In the presence of Dox, the total number of DO11 cells in both BALB/c and Ii-TGO hosts increased, but remarkably, the number of DO11 Tregs increased by about 2-fold in only the Ii-TGO hosts (Figure 3A-C). Moreover, in the presence of Dox, the percentage of DO11 cells expressing both the activation marker CD69 and the lymph node homing receptor CD62L increased in both BALB/c hosts and Ii-TGO hosts. Interestingly, almost half of this CD69^+^CD62L^+^ population were also CD25^+^Foxp3^+^ in Ii-TGO hosts on Dox, but <10% of CD69^+^CD62L^+^ T cells in BALB/c hosts on Dox were Tregs (Figure 3D-E). Both CD69 and CD62L expression by Tregs has been associated with more effective anti-inflammatory function in other models of sterile inflammation (32-34).

**Figure 3.**
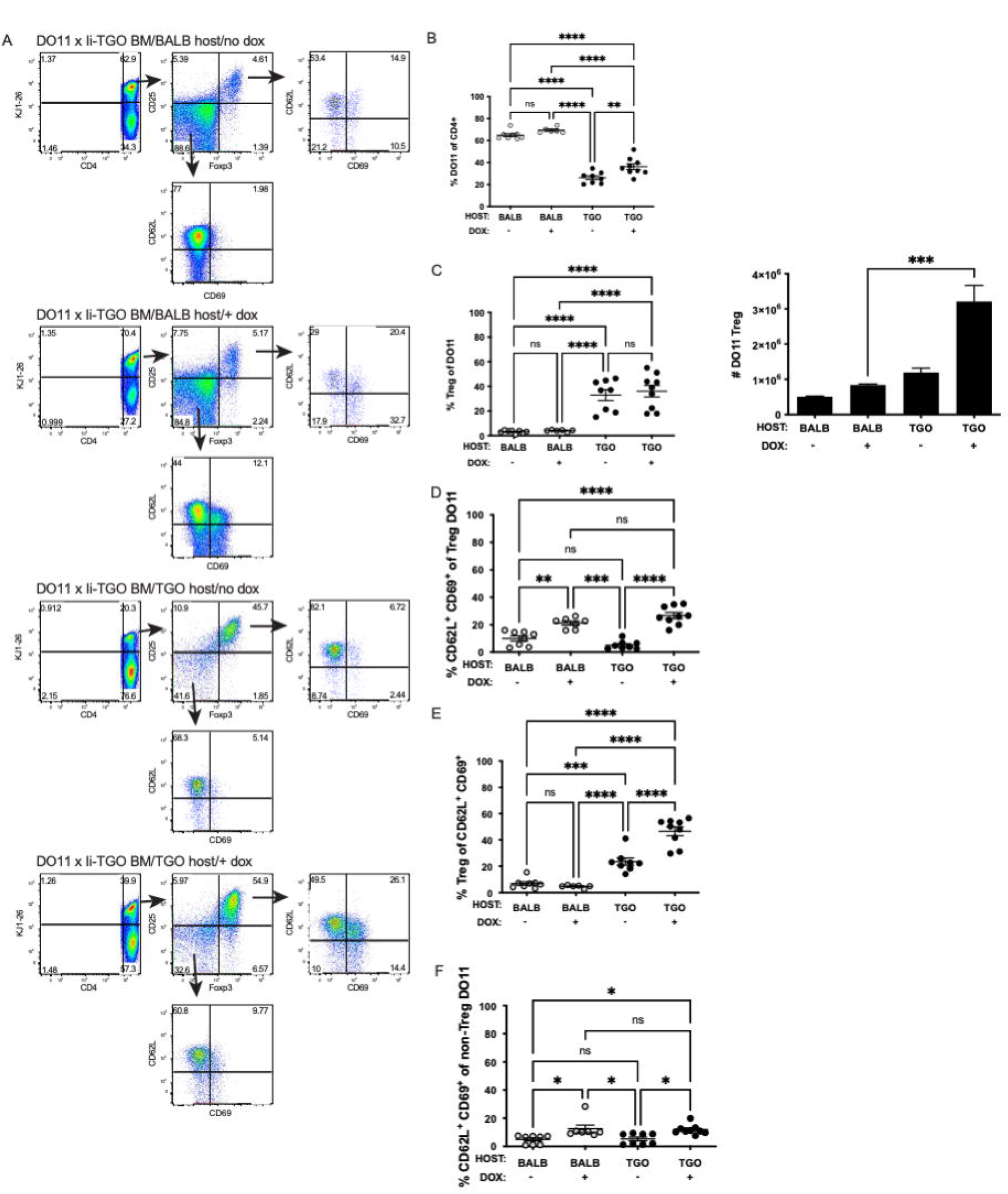
Self-Ag presentation by radioresistant APCs drives activation and elaboration of SLO-tethered Tregs. (A) DO11 × Ii-TGO bone marrow was used to reconstitute lethally irradiated BALB/c and TGO hosts. At 12 weeks, the mice were either maintained on normal chow or placed on Dox chow. The mice were sacrificed 7 days later, and spleen cell suspensions were stained and analyzed by flow cytometry. Representative plots of the four groups are provided. (B-F) Summary of combined data from two independent experiments; line indicates the SEM. (B) Frequency of DO11 (KJ1-26+) cells within the CD4 gate. (C) Frequency of Tregs (CD25+Foxp3+) within the KJ1-26+ gate. (D) Frequency of CD62L+ CD69+ cells within the KJ1-26+ Treg gate. (E) Frequency of Tregs within the KJ1-26+ CD62L+ CD69+ gate. (F) Frequency of CD62L+ CD69+ cells within the KJ1-26+ non-Treg gate. Student’s *t*-test was used for statistical analysis.

To determine whether radioresistant lymph node stroma could (1) expand Tregs and (2) sustain or enhance CD62L expression on activated Tregs, we co-cultured splenic CD11c^+^ or CD45^-^ cells with CD4^+^ cells from DO11-TGO mice. LNSC and splenic stromal cells induced Treg proliferation in response to OVA peptide. CD62L expression was high in LNSC/T cell co-cultures, while CD11c^+^ APC/T cell co-cultures led to downregulation of CD62L on Tregs (Figure 4A). These data show that SLO stromal cells induce Treg proliferation while poising Tregs to remain in secondary lymphoid organs. Strikingly, stroma-directed reinforcement of CD62L expression was robust in the Treg population, but not in the effector cell population of the same cultures (Figure 4B). Thus, we have identified a previously unrecognized player in dominant systemic tolerance, the SLO stromal cell, which both activates and poises Foxp3^+^ Tregs to suppress activation of unprimed autoreactive T and B cells in the spleen or lymph nodes through sustained expression of CD62L following activation.

**Figure 4.**
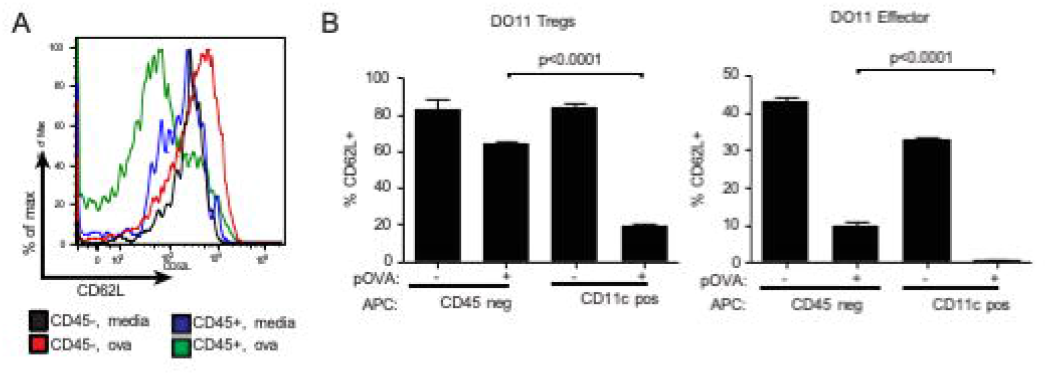
Cognate Ag presentation by LNSC reinforces Treg capacity for lymph node homing, but not effector T cells. (A) Histogram depicting log^10^ fluorescence of CD62L on DO11 Tregs after culture with magnetically separated LNSC. (B) CD45- stromal cells or CD11c+ cells purified from pooled lymph nodes of Ii-TGO mice were cultured with CD4+ T cells from DO11-Foxp3^eGFP^ x Ii-TGO mice. Plot shows the frequency of CD62L^high^ cells within the KJ1-26+ eGFP+ gate. Graph shows the frequency of CD62L^high^ cells within the KJ1-26+ eGFP-gate The results shown are pooled from three experiments.

## Discussion

LNSC regulation of immune responses has been described (19, 20, 23, 24); however, the programming of LN-homing Tregs by LNSC is novel. The data described here indicate that autoantigen-presenting radioresistant host cells play a critical role in limiting the expansion of autoreactive T cells and driving naïve autoreactive T cells into the Treg compartment. Taken together, these data indicate that a radioresistant peripheral host cell population bearing self-Ag most effectively activates and expands Tregs which become “tethered” to secondary lymphoid organs where they could potentially suppress autoreactive B cell activation and ANA production.

We have shown that T cell activation is an important checkpoint in the prevention of autoantibody production, that autoreactive T cell activation is sufficient for driving autoantibody production by autoreactive B cells, and that antigen-specific Tregs most effectively control autoantibody responses. Notably, our experiments examine the behavior of normal B cell repertoire, presumably including a small frequency of autoreactive B cells, which are availed of cognate interactions with T cells. We show that even with Ag-specific interactions, naïve T cells could not drive ANA production from endogenous autoreactive B cells. DO11 T cells from naïve mice divide in Ii-TGO mice, but did not stimulate ANA production under conditions where activated DO11 T cells could stimulate ANAs.

TGO expression by rAPCs led to activation of DO11 Tregs (using CD69 expression as criteria for activation) while maintaining expression of CD62L, which is usually downregulated after priming of naïve T cells (although it can later be upregulated) (Figure 3, 4). CD62L^+^ Tregs are potent suppressors of graft-versus-host disease and in a model of type 1 diabetes, most likely because they home preferentially to lymph nodes and can suppress primary immune responses (32-34). We determined that expression of self-antigen by radioresistant host cells could drive Treg differentiation. Upregulation of CD62L on Tregs in these cultures suggests that the SLO stromal cells can activate and poise Foxp3^+^ Tregs to suppress activation of naïve autoreactive T and B cells in spleen or lymph node through sustained expression of CD62L following activation. This phenomenon appears to be Treg specific, and the large presence of non-Tregs in CD4-purified cell/stromal cell cultures slightly abrogated CD69 maintenance on Tregs (Figure 3, 4). Future directions include determination of the specific stromal-derived signals that promote LN-homing Treg and studies in mice with CD62L deficient Tregs.

## Materials and Methods

### Mice

BALB/c, BALB/c.Foxp3^eGFP^, and DO11.10 mice were purchased from the Jackson Laboratory and maintained in our colony for at least 1 week prior to use. DO11.10 Rag2-/-mice were bred in-house after purchase from Taconic. See Mande et al for additional description of the generation of Ii-TGO mice (35). In some experiments, mice were fed *ad lib* with Dox chow (200mg doxycycline/kg of chow; Bioserv). All mouse experiments were approved by the Institutional Animal Care and Use Committee of the University of Massachusetts Medical School. In all experiments, the mice were age-matched within 2 weeks of each date of birth.

### Radiation chimeras

Mice were irradiated using a Cesium-137 irradiator in rotating containers. For adoptive transfers of mature T cells, mice were dosed with 400 rads and injected the following day. For lethal irradiation prior to bone marrow transfer, mice were dosed with 800 rads and reconstituted with 10^7^ T-depleted bone marrow cells the following day.

### Flow cytometry

Single cell suspensions were surface or intracellularly stained with combinations the following antibodies: I-A/I-E-FITC, CD3 (clone 145-2C11), CD4 (clone RM4-5), CD8α (clone 53-6.7), CD25-PerCP-Cy5.5 (clone PC61.5), CD69-efluor450, CD62L-APC-efluor780, B220, Foxp3-PE (clone FJK-16s) all from eBioscience, and CD45-APC (Miltenyi). For Foxp3 staining, cells were fixed and permeabilized using the Foxp3 transcription factor staining buffer kit (eBioscience) according to the manufacturer’s instructions.

### In vitro T cell differentiation and transfer

CD4+ T cells were purified from DO11 lymph node and spleen cell suspensions by positive selection (BD iMag CD4 beads; manufacturer’s instructions). Cells were cultured in complete RPMI medium 1640 (RPMI; Life Technologies, containing 10% FCS (Sigma), 10 µM 2-mercaptoethanol, 100 units/ml penicillin, 0.1 mg/ml streptomycin, and 0.3 mg/ml glutamine (GIBCO) and stimulated with irradiated BALB/c spleen (3,000 rads; 1:5 ratio of T cells to irradiated spleen) for 48–72 h in the presence of OVA peptide (pOVA,1µg/ml; Genscript), X63.Il4 supernatant (1:4), and X63.Il2 supernatant (2 drops per 5 ml), and split as needed. On day 7–9, cultures were re-stimulated with irradiated BALB/c spleen (3,000 rads) for 48–72 h in the presence of pOVA; cells (5 × 10^6^) cells were injected per mouse i.v. via tail vein. For some experiments, naïve T cells were labeled with CFSE according to the manufacturer’s (Invitrogen) recommendations, prior to i.v. injection.

### HEp2 ANA assay

Sera were diluted 1:50 in PBS were assayed as described previously (36)

### Stromal cell and CD11c+ cell isolation

Lymph nodes and spleens were placed in a Petri dish containing pre-warmed RPMI with 2% FBS containing 0.5 mg/ml dispase and 0.5 mg/ml collagenase. The tissues were gently disrupted mechanically with frosted glass slides, and then incubated for 15 minutes at 37°C. A sterile P1000 pipet tip was cut with an autoclaved razor to increase the size of the opening. This large-bore tip was used to disrupt the tissues by pipetting. The enzyme cocktail was removed by pipetting and the tissue was washed four times with RPMI. Fresh enzyme cocktail was added to the remaining solid tissue, and this process was repeated twice more. After four washes, single cell suspensions of lymph nodes and spleen were used for CD11c positive selection or CD45 depletion (Miltenyi; according to manufacturer’s instructions).

### Statistical methods

Data analysis was performed using Prism (GraphPad) software. *p* values were calculated using unpaired two-tailed Student’s t-test or 2-way ANOVA, as indicated in figure legends. In all figures, ns = not significant (*p*>0.05), *, *p*≤0.05, **, *p*≤0.01, ***, *p*≤0.001.

## ACKNOWLEDGEMENTS

This work was supported by National Institutes of Health Grants AR055634 (to AM-R). TYB was supported by T32 AI095213. The authors would like to thank Tara E. Robidoux for superb animal care and technical assistance and Karen Lee MD, PhD for early development of the original Ii-TGO model.

## Figure Legends

**Supplemental Figure 1. Autoreactive T cell activation is required for autoantibody production**. (A) Schematic of the breeding strategy for generating Ii-TGO mice. (B) 5 × 10^6^ naïve DO11 or DO11 Th2 cells were adoptively transferred into sublethally irradiated (400 rads) Ii-TGO mice fed normal or dox chow. Mice were cheek-bled at 6 weeks and serum was analyzed by HEp2 ANA assay. (C) The total antibody concentration in Ii-TGO sera post T cell transfer was quantified by isotype specific ELISAs. Ii-TGO mice were irradiated (400 rads), placed on Dox chow, and i.v. injected with DO11 Th2 cells.

