## Supplementary figures and images for "Secondary lymphoid organ stroma activate and elaborate regulatory T cells to suppress autoantibody production in a novel model of systemic autoimmunity"

### Supplemental Figure 1

#### Supplemental Figure 1

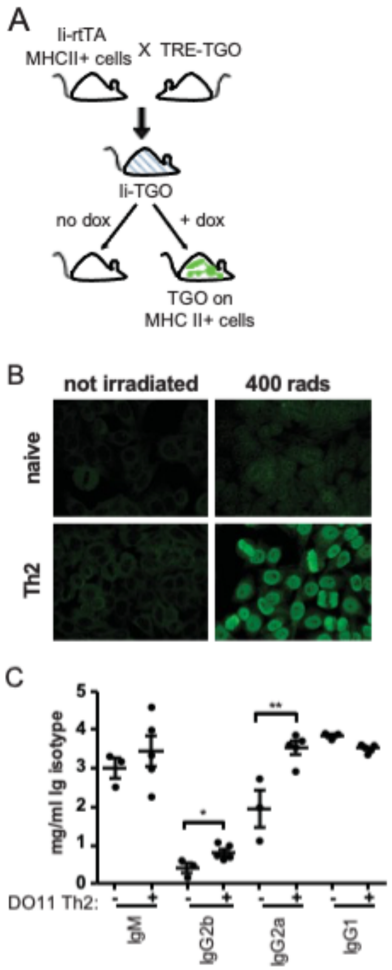
